# Rapid Epistatic Mixed Model Association Studies by Controlling Multiple Polygenic Effects

**DOI:** 10.1101/2020.03.05.976498

**Authors:** Dan Wang, Hui Tang, Jian-Feng Liu, Shizhong Xu, Qin Zhang, Chao Ning

## Abstract

**Summary:** We have developed a rapid mixed model algorithm for exhaustive genome-wide epistatic association analysis by controlling multiple polygenic effects. Our model can simultaneously handle additive by additive epistasis, dominance by dominance epistasis and additive by dominance epistasis, and account for intrasubject fluctuations due to individuals with repeated records. Furthermore, we suggest a simple but efficient approximate algorithm, which allows examination of all pairwise interactions in a remarkably fast manner of linear with population size. Application to publicly available yeast and human data has showed that our mixed model-based method has similar performance with simple linear model-based Plink on computational efficiency. It took less than 40 hours for the pairwise analysis of 5,000 individuals genotyped with roughly 350,000 SNPs with five threads on Intel Xeon E5 2.6GHz CPU.

**Availability and implementation:** Source codes are freely available at https://github.com/chaoning/GMAT.

## 1. Introduction

Epistasis is defined as the deviation of the combined allele effect at two or more loci from the sum of their individual effects (Fisher, 1918). It is considered a promising approach to understanding genetic causes of complex traits (Mackay and Moore, 2014; Phillips, 2008; Upton, et al., 2016).

Linear mixed models have been widely used in genome-wide association studies (GWAS) due to its advantage in correcting environmental factors, controlling population stratification and accounting for cryptic relatedness between individuals (Kang, et al., 2008; Lippert, et al., 2011; Zhou and Stephens, 2012). However, computational inefficiency hindered the application of linear mixed models in performing exhaustive epistatic association studies with the increases of sample size and marker density. For example, Lippert, et al. (2013) applied linear mixed models to the exhaustive epistatic SNP association studies across seven traits for 14,925 individuals genotyped with 356,441 SNPs. It would take 950 computer years, with a wall-clock time of 13 days using 28,000 cores.

To address the issues, we developed a rapid epistatic mixed model association study (REMMA) in previous studies (Ning, et al., 2018). REMMA has an advantage over existing linear mixed models in terms of high computational efficiency, reduced type I error and high QTN detection power. However, the time complexity of computation is still *O*(*n*^2^), where *n* is the population size. Furthermore, REMMA only focuses on the additive by additive epistatic analysis and cannot deal with study population consisting of individuals with repeated records. We report here a REMMA eXpedited (REMMAX) version of the method. REMMAX can simultaneously deal with the association studies for additive by additive epistasis, dominance by dominance epistasis and additive by dominance epistasis by incorporating multiple polygenic effects as background control. Furthermore, individual-specific residual effect has been included in the model to control intrasubject fluctuations due to individuals with repeated records. To improve the computational efficiency, we also propose an approximate REMMAX algorithm, which first implements an approximate statistical test with computational complexity of *O*(*n*) to filter out many nonsignificant interactions and then employs the exact Wald Chi-square test to test the interaction effects of selected SNP pairs. Experiments on yeast (Bloom, et al., 2015) and WTCCC (Wellcome Trust Case Control, 2007) data sets show that REMMAX is an efficient method to perform the exhaustive epistatic analysis.

## 2. Methods

### 2.1 REMMAX Model

A linear mixed model including additive and non-additive SNP effects can be written as

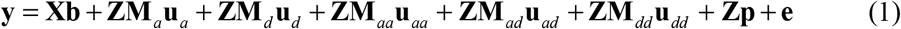

where **y** is a vector of *N* observations for *n* individuals (including repeated measurements). Below is the description of the model effects and their design matrices. Vector **b** is a collection of fixed effects. Vectors **u**_*a*_, **u**_*d*_, **u**_*aa*_, **u**_*ad*_ and **u**_*dd*_ are additive, dominance, additive by additive, additive by dominance and dominance by dominance SNP effects, respectively. Vector **p** is an array of individual-specific residual effects. Vector **e** is a collection of random residual errors. **X** is a design matrix for the fixed effects. **Z** is a design matrix for all the random model effects, serving as connection between the model effects and observed data points in the response variable. **M**_*a*_, **M**_*d*_, **M**_*aa*_, **M**_*ad*_ and **M**_*dd*_ are additive, dominance, additive by additive, additive by dominance and dominance by dominance SNP matrixes, respectively. They are formalized as

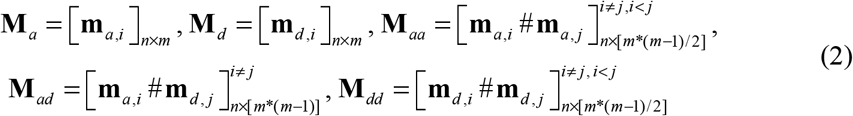

Here, *m* is the number of SNPs; # is Hadamard (element-wise) product. **m**_*a,i*_ and **m**_*d,i*_ are additive and dominance SNP vectors for the *i*th SNP with elements defined as:

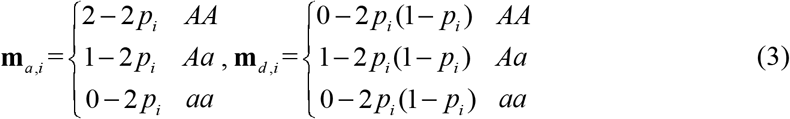

Where *p_i_* is the allele frequency of allele A for the *i*th SNP. Below are the assumed normal distributions for the random effects:

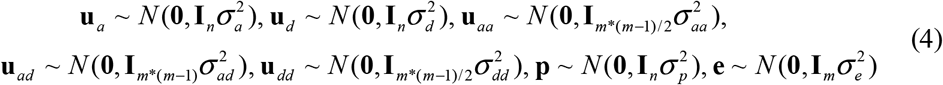

Here, 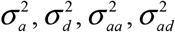 and 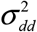 are additive, dominance, additive by additive, additive by dominance and dominance by dominance variances; **I**_*_ is identity matrix; Finally, 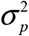 and 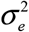 are individual-specific error and random error variances, respectively. The variances can be estimated by maximizing the following restricted log-likelihood function with efficient combined expectation maximization (EM) and average information (AI) algorithm (Ning, et al., 2019).

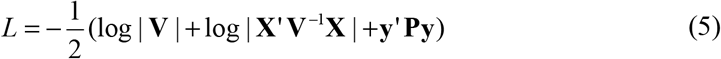

Where

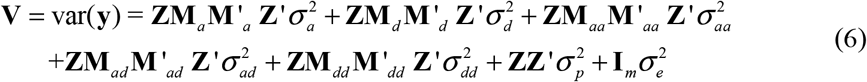

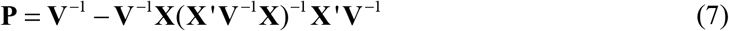

However, the calculation of **V** is inefficient due to high-dimensional **M**_*aa*_, **M**_*ad*_ and **M**_*dd*_. According to previous studies (Ning, et al., 2018; Su, et al., 2012; VanRaden, 2008; Xu, 2013), 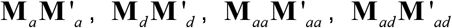 and 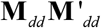 can be used to respectively reflect the additive, dominance, additive by additive, additive by dominance and dominance by dominance genomic relationship among individuals with the following equation.

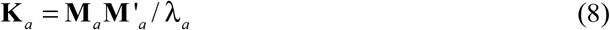

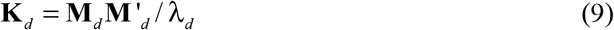

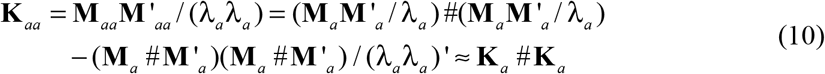

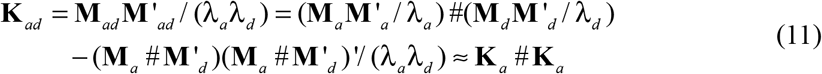

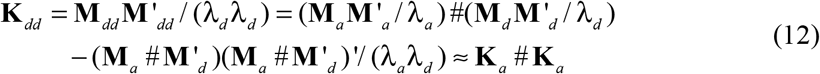

Where, λ_*a*_ = ∑2*p*_*i*_(1 − *p*_*i*_), λ_*d*_ = ∑ 2*p*_*i*_(1 − *p*_*i*_)(1 − 2*p*_*i*_(1 − *p*_*i*_)). Apply equation (8–12) to (6), **V** can be efficiently calculated with time complexity of *O*(*n*^2^*m*) in the following equations

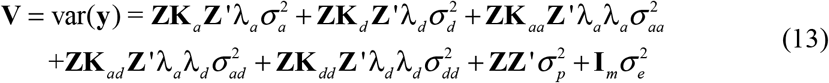

Following Henderson (1975), the random SNP effects can be estimated with the following equations:

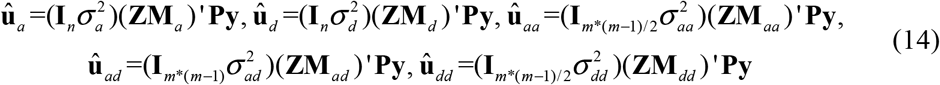

The corresponding variances are

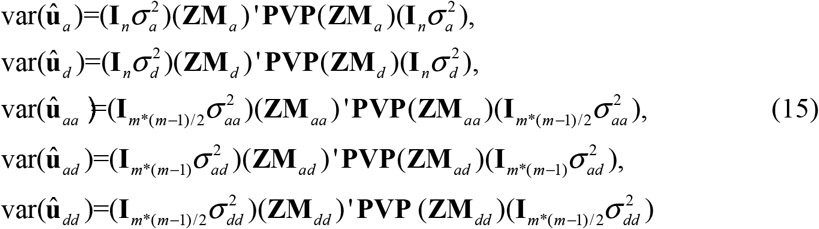

For the *i*th and *j*th SNPs, the estimated SNP effects and corresponding variances are

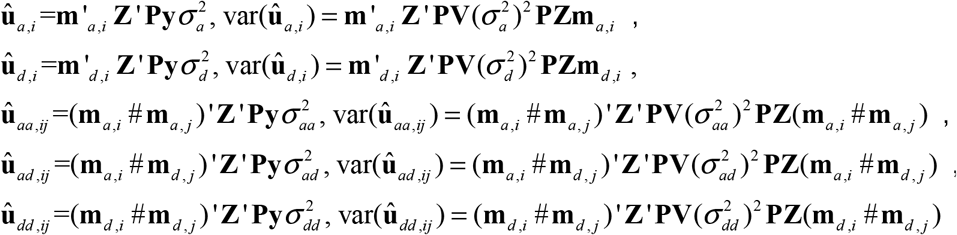

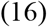

The Wald Chi-squared test is defined as:

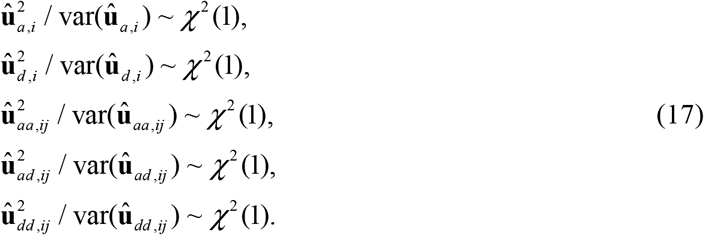

Note that **Z**’**Py** and **Z**’**P** var(**y**)**PZ** can be precalculated. The SNP effects are calculated by vector-vector multiplication whose computational complexity is *O*(*N*), while the variances of estimated effects are calculated by vector-matrix-vector multiplication whose computational complexity is *O*(*N*^2^).

### 2.2 Approximate algorithm

To reduce the time complexity, an efficient algorithm with the approximate Wald Chi-squared statistical test is implemented as following

Step 1: Randomly select *k* (e.g. 100,000) SNP pairs and calculate their epistatic effects and the corresponding variances. The median of variances is defined as var(*û*)_*median*_.

This step has time complexity of *O*(*kN*^2^)

Step 2: Scan the genome-wide epistatic effects and calculate the approximate Wald Chi-squared statistical test with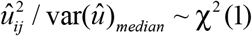. This step has time complexity of *O*(*Tm*), where *T* is the total number of pairwise test. *T* is *m*(*m*−1)/2 for additive by additive and dominance by dominance epistatic test, and is *m*(*m*−1) for additive by dominance epistatic test, where *m* is the number of SNPs.

Step 3: Select top significant SNP pairs based on the approximate Wald Chi-squared statistical test using a *p*-value threshold (e.g. *p*-cut = 10^−5^). This step has a time complexity of *O*(*rN*^2^), where *r* is the number of top significant SNP pairs.

For population with over hundred thousand SNPs, *kN* and *rN* have similar size to *T*. Therefore, for each SNP pair epistatic test, the time complexity can be approximated by [*O*(*km*^2^)+*O*(*Tm*)+ *O*(*rm*^2^)]/*T*≈*O*(*m*).

### 2.3 Data Analysis

We applied the REMMAX method to the yeast data (Bloom, et al., 2015) and the WTCCC data (Wellcome Trust Case Control, 2007). The yeast data contained 4,390 BY×BM segregants genotyped with 28,220 markers (no heterozygous genotype). Twenty quantitative traits with repeated measurements were analyzed with the following model

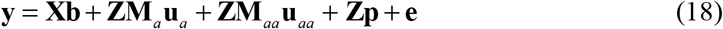

The rheumatoid arthritis (RA) and type-I diabetes (T1D) traits were analyzed for the WTCCC data. After quality control of genotypes following Ning, et al. (2018), 4,961 individuals genotyped by 353,859 SNPs for RA and 4,962 individuals genotyped by 353,751 for T1D were remained for subsequent analysis. As no individuals has repeated records, **p** is removed from model and **Z** is the identity matrix. Then, the data was analyzed with the following model

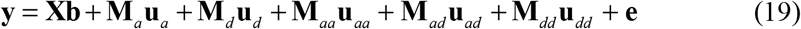

To observe whether diffident levels of polygenic effects will affect the additive by additive epistatic test, the following models were also included to analyze WTCCC data.

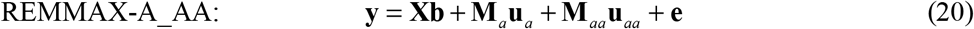

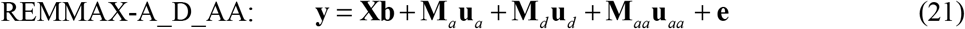

Here, the REMMAX-A_AA model with only additive and additive by additive SNP effects is the same to REMMA method of our previous studies (Ning, et al., 2018).

## 3. Results

We observed high Pearson correlation coefficients of additive by additive epistatic *P* values from models with diffident levels of polygenic effects (**Figure 1**). It indicates that if we focus on additive by additive epistatic test, previous REMMA method can also work well.

**Figure 1.**
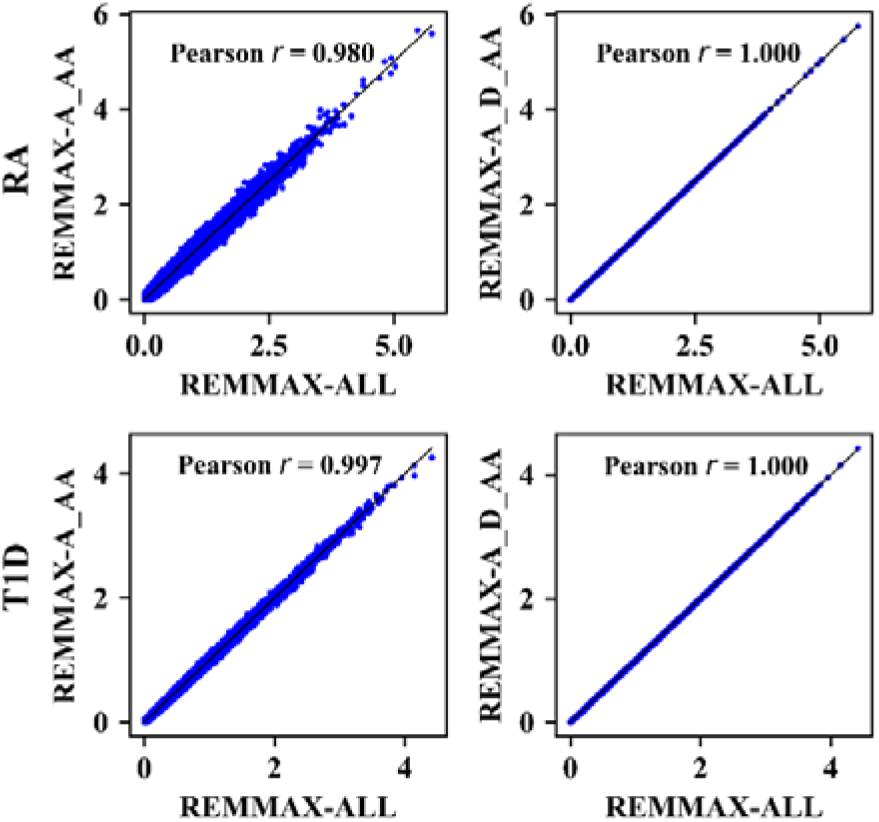
Comparisons of additive by additive epistatic *P* values from models with diffident levels of polygenic effects in the analysis of RA and T1D traits of WTCCC data. REMMAX-A_AA, REMMAX-A_D_AA and REMMAX-ALL refers to equation (20), (21) and (19), respectively. Random 100,000 SNP pairs were selected for comparison.

In order to assess whether REMMAX can control type I error well, we randomly shuffled analysis SNP across individuals at each locus, which can purposely destroy the association of the phenotypes with the scanned SNP. Under the expectation that random SNP pairs are unlinked to polymorphisms controlling these traits, the cumulative *P*-value distribution follows a uniform distribution of *U*(0, 1). The **Figure 2 and Supplementary Figure 1** show that the type I errors are well controlled by the exact Wald Chi-squared statistical tests, but are not controlled by approximate Wald Chi-squared statistical tests in some cases.

**Figure 2.**
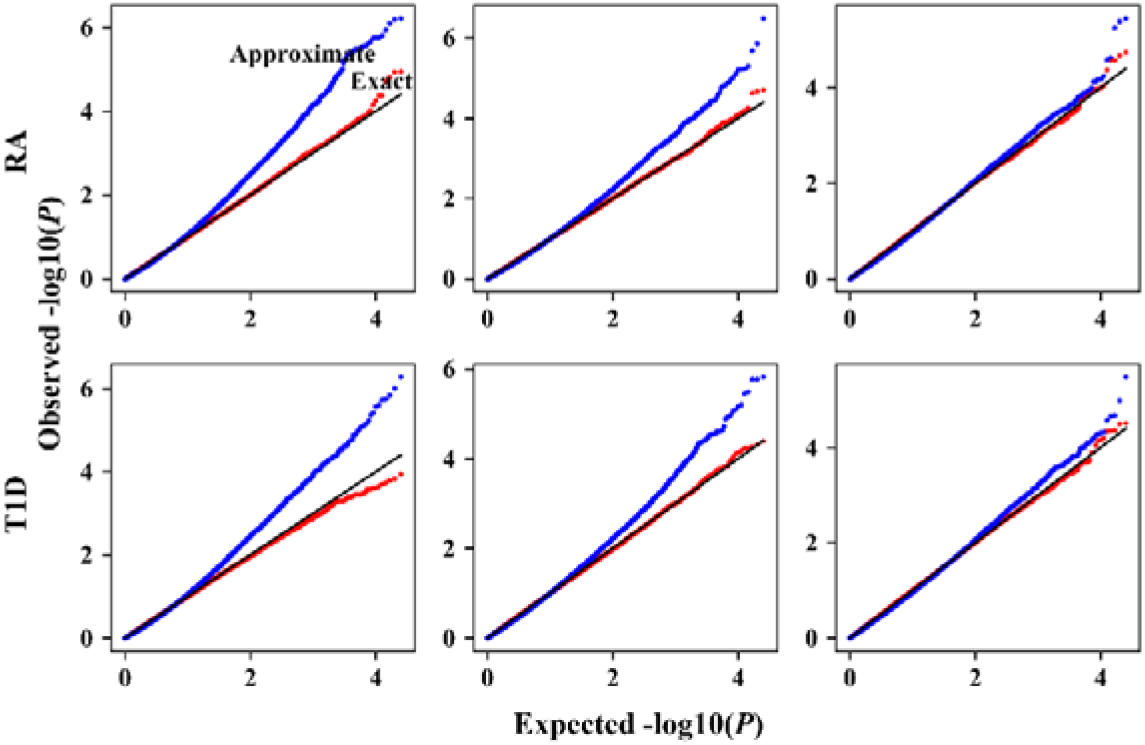
The quantile-quantile plots of exact (red) and approximate (blue) Wald Chi-squared statistical tests in testing additive by additive (left), additive by dominance (middle) and dominance by dominance (right) epistatic SNP effects. We random selected 100,000 SNP pairs to estimate the null distribution.

Then, we compared the *P*-values obtained by the approximate and exact Wald Chi-squared statistical tests in the real data analysis of the yeast and WTCCC data. A high correlation coefficient was observed in the WTCCC data for the RA and T1D traits (**Figure 3**). The Pearson correlation coefficients between the approximate and exact test are all very closed to unity for the yeast data (**Supplementary Figure 2**). Integrating the results of type I error and high correlation coefficients of approximate and exact Wald Chi-squared statistical tests, the approximate REMMAX method, which first implements an approximate statistical test to filter out many nonsignificant interactions and then employs the exact Wald Chi-square test to confirm the true significant SNP pairs, is theoretically feasible to genome-wide epistatic association studies.

**Figure 3.**
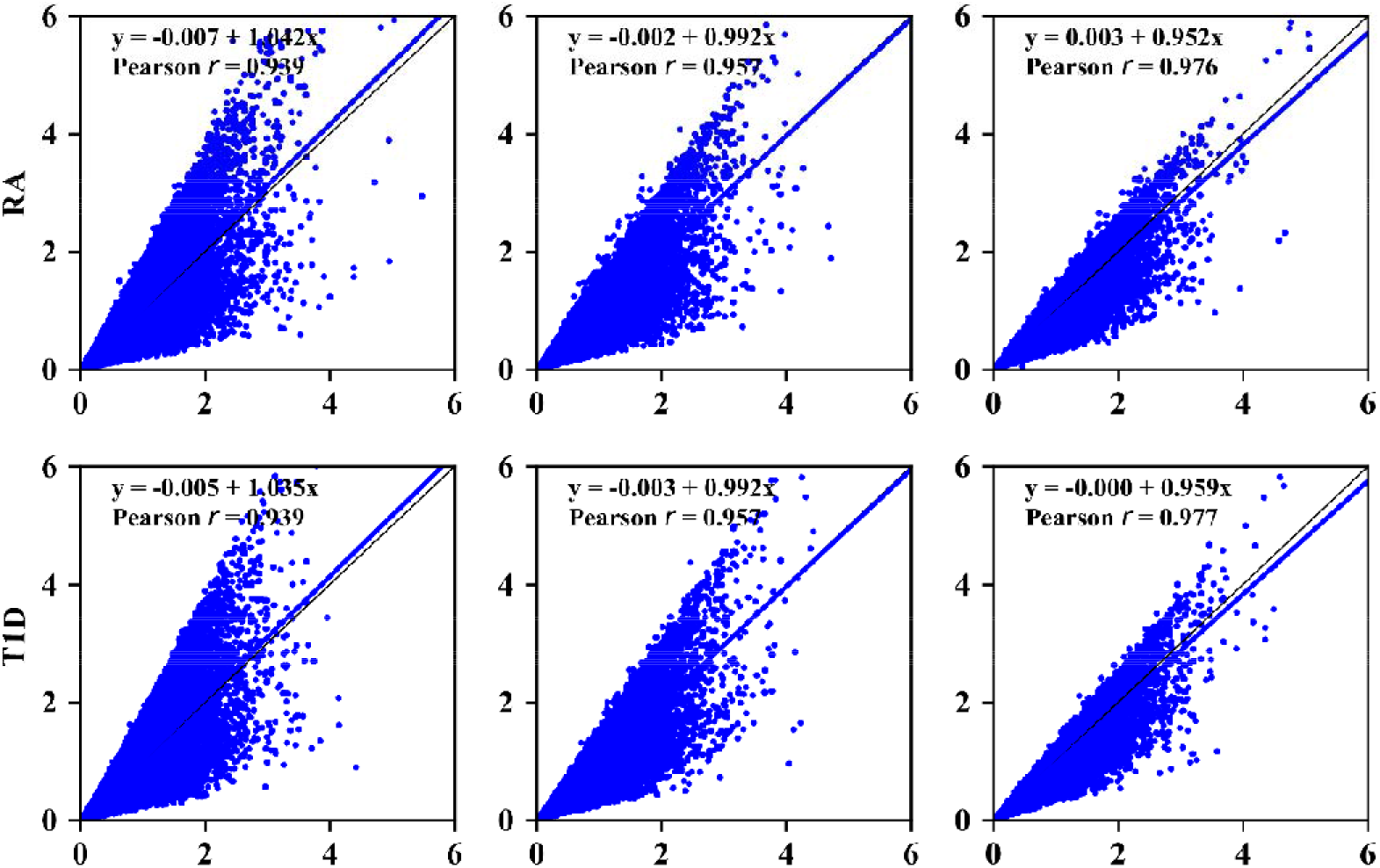
Comparisons of *P* values obtained by exact (X axis) and approximate (Y axis) REAMMX statistical tests for additive by additive (left), additive by dominance (middle) and dominance by dominance epistasis (right) in the WTCCC data. The blue lines mean lines of best fit, while black lines mean y=x.

In the analysis of 20 traits in yeast data, we located 665,195 significant SNP pairs by approximate REMMAX after Bonferroni correction. Only nine SNP pairs were missed compared to exact REMMAX (**Supplementary Table 1**). As it is very time consuming to apply the exact REMMAX to the genome-wide pairwise analysis for the WTCCC data, we only applied the approximate REMMAX method to the WTCCC data. It took no more than 40 hours for each the pairwise analysis with five threads on Intel Xeon E5 2.6GHz CPU. The predicted time is one year for exact REMMAX method under the same condition. For the RA trait in the WTCCC data, the approximate REMMAX method located 978 significant SNP pairs (198 for additive by additive, 606 for additive by dominance and 174 for dominance by dominance), and 664 of them (67.9%) were in the MHC region (chr6:28,510,120-33,480,577). For T1D trait, we also observed most of the significant SNP pairs (2808/2836 for additive by additive, 1404/2303 for additive by dominance and 3555/3619 for dominance by dominance) were in the MHC region. These results are consistent with previous studies (Ning, et al., 2018; Wan, et al., 2010). For comparison, we applied the exact REMMAX method to pairwise analysis of the chromosome 6. Taking the results of exact REMMAX as the gold standard, approximate REMMAX located most of the significant SNP pairs (94.6% for RA and 99.8% for T1D).

We further compared the time complexities and the actual running times of approximate REMMA and exact REMMAX compared to simple linear model implemented by Plink (Purcell, et al., 2007) (**Table 1**). Simple linear model is the most efficient with time complexity of *O*(*N*). As *k* and *r* are much smaller than *T*, the time complexity for REMMAX is nearly linear with *N*. For both the yeast and human data, approximate REMMAX was able to finish the exhaustive analysis within a tolerable time frame over a hundred times faster than exact REMMAX.

**Table 1.** Time complexities and computing times of simple linear model, exact REMMAX and approximate REMMAX for exhaustive genome-wide epistatic studies.

| Method | Time complexity | Computing time |  |
| --- | --- | --- | --- |
|  |  | Yeast | WTCCC |
| Simple linear model | $O(N)$ | 5.4min | 4.7min |
| exact REMMAX | $O(N^2)$ | 32.79h | 38h |
| approximate REMMAX | $O(kN^2) + O(TN) + O(rN^2)$ | 6.5min | 7.2min |

*Note*: All computing was performed with five threads on Intel Xeon E5 2.6GHz CPU.

*N* is the number of observations for all individuals; *k* is the number of randomly selected SNP pairs; *T* is the total number of pair-wise tests; *r* is the remaining number of SNP pairs filtered by the approximate Wald Chi-squared statistical test. The simple linear model is implemented by Plink. As Plink cannot deal with study population of individuals with repeated records, we used the average values as the response variable for Plink in the yeast data. We only applied the additive by additive epistatic analysis to the chromosome 6 in the WTCCC data, as Plink can only deal with additive by additive epistatic analysis and it is time consuming for exact REMMAX to perform the exhaustive analysis.

## 4. Discussion

In this study, we proposed a rapid epistatic mixed model association studies method, REMMAX. REMMAX can handle the additive by additive epistasis, additive by dominance epistasis and dominance by dominance epistasis, and individual-specific residual effect has been included in the model to analyze data with repeated measurements. Furthermore, we build an approximate Wald Chi-squared statistical test for REMMAX to expedite the exhaustive genome-wide epistatic association studies. The time complexity is nearly linear with the population size. Experiments on yeast and WTCCC data sets show that REMMAX is an efficient method to help understand genetic constitutions of complex traits.

Our previous REMMA (Ning, et al., 2018) method is the special case of REMMAX, where only genome-wide additive and additive by additive SNP effects are included. When we analyze the cross-sectional data (one record per individual), the individual-specific residual effect is same to random residual and should be removed from the model. In the study, we have learned that diffident levels of polygenic effects have little influence on the additive by additive epistatic association studies. Therefore, the REMMA method is also preferred when we focus on the additive by additive analysis of cross-sectional data. Additionally, in the estimation of variances of estimated SNP effects, REMMA has to calculate the inverse of huge mixed model equation which is affected by the number of random effects included in linear mixed model. For REMMAX, the computational bottleneck is to calculate the inverse of phenotypic covariance matrix (**V**), whose dimension is always of *N* (*N* is equal to *n* for cross-sectional data) regardless of the number of random effects. For example, the dimension of mixed model equation and phenotypic covariance matrix is 6*n* and *N*, respectively. Therefore, the amount of computation and memory is reduced to about 1/27, thus the computation time and memory consumption is shortened significantly.

## Acknowledgments

This study makes use of data generated by the Wellcome Trust Case-Control Consortium. A full list of the investigators who contributed to the generation of the data is available from www.wtccc.org.uk. Funding for the project was provided by the Wellcome Trust under award 076113 and 085475.

