## Supplementary Information for "Rapid Epistatic Mixed Model Association Studies by Controlling Multiple Polygenic Effects"

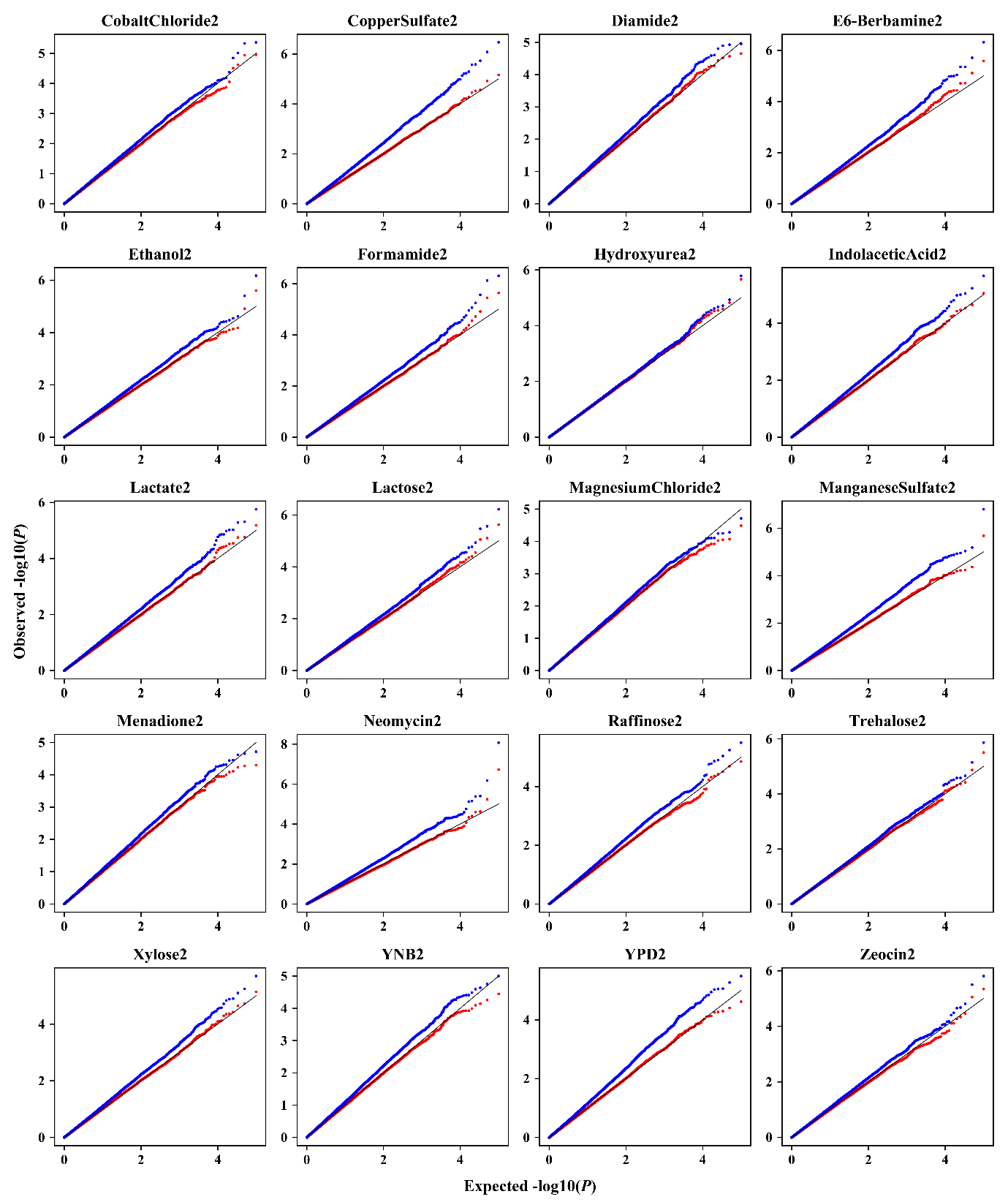

**Supplementary Figure 1.** The quantile-quantile plots of exact (red) and approximate (blue) Wald Chi-squared statistical tests in testing additive by additive epistatic SNP effects for the 20 traits in the yeast data. We random selected 100,000 SNP pairs to estimate the null distribution.

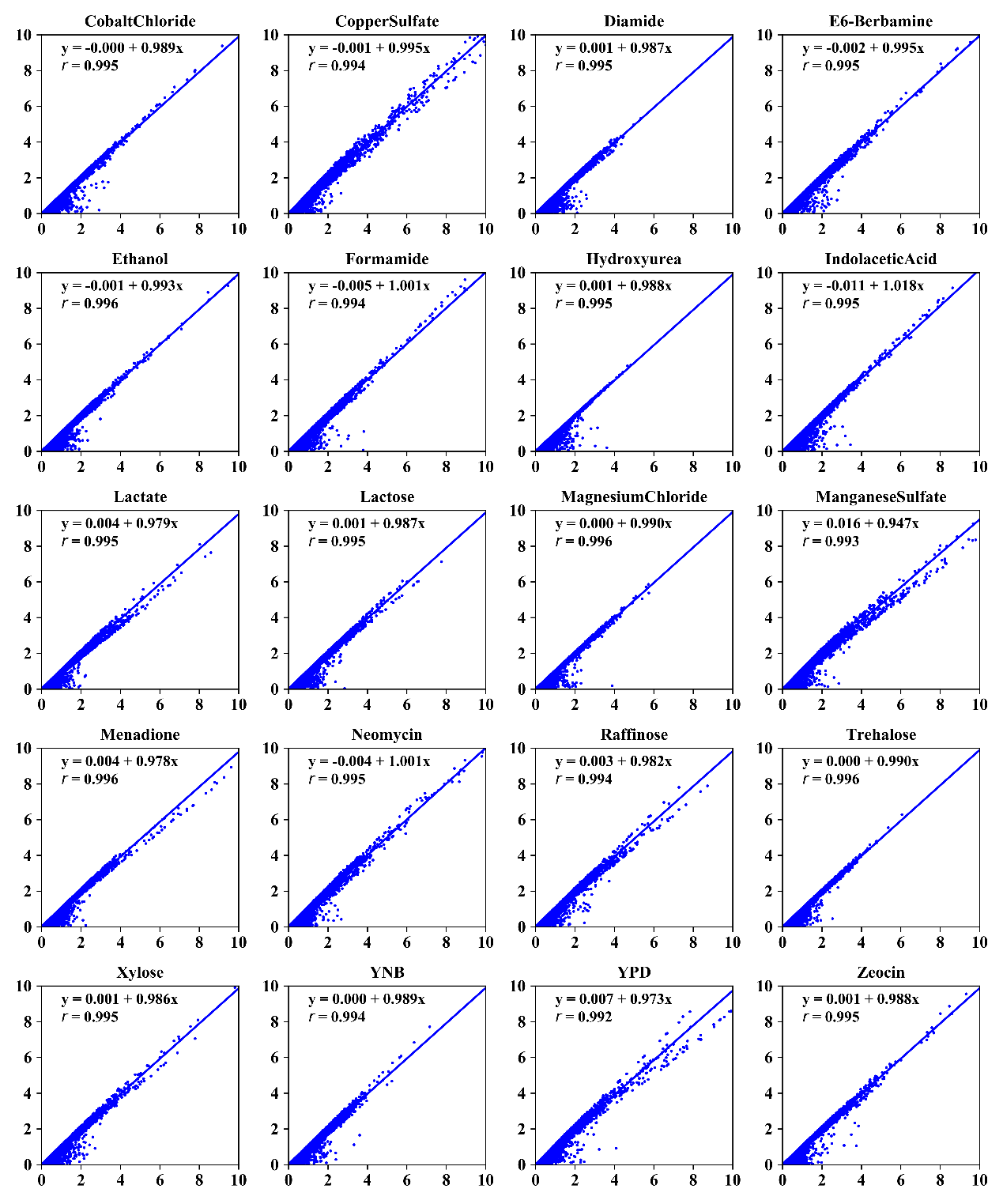

**Supplementary Figure 2.** Comparisons of *P* values obtained by exact (X axis) and approximate (Y axis) REMMAX statistical tests for the 20 traits in the yeast data. The blue lines mean lines of best fit, while black lines mean y=x.

**Supplementary Table 1.** The number of significant SNP pairs located by exact REMMAX and approximate REMMAX in the yeast data.

| Trait | exact REMMAX | approximate REMMAX |
| --- | --- | --- |
| CopperSulfate | 126,613 | 126,609 |
| CobaltChloride | 2,174 | 2,174 |
| Menadione | 39,482 | 39,482 |
| Zeocin | 4,274 | 4,273 |
| Ethanol | 52,424 | 52,424 |
| Neomycin | 69,155 | 69,155 |
| YPD | 44,285 | 44,285 |
| Trehalose | 0 | 0 |
| Xylose | 3,625 | 3,625 |
| MagnesiumChloride | 0 | 0 |
| Formamide | 72,050 | 72,050 |
| YNB | 4,662 | 4,662 |
| Lactate | 22 | 21 |
| ManganeseSulfate | 131,220 | 131,220 |
| IndolaceticAcid | 83,248 | 83,248 |
| Lactose | 2 | 0 |
| Raffinose | 3 | 2 |
| Hydroxyurea | 0 | 0 |
| E6-Berbamine | 31,965 | 31,965 |
| Diamide | 0 | 0 |
| sum | 665,204 | 665,195 |
